# Fibrotic human lung extracellular matrix as a disease-specific substrate for 3D *in-vitro* models of pulmonary fibrosis

**DOI:** 10.1101/833913

**Authors:** Igal Germanguz, Evelyn Aranda, Jennifer C. Xiong, Natalia Kissel, Alexandra Nichols, Eddie Gadee, John D. O’Neill

**Affiliations:** Xylyx Bio, Inc., 760 Parkside Avenue, Brooklyn, New York 11226, USA

**Keywords:** 3D cell culture, drug testing, extracellular matrix, idiopathic pulmonary fibrosis, *in-vitro* models, lung disease, lung fibroblasts, scaffolds

## Abstract

Idiopathic pulmonary fibrosis (IPF) is an irreversible and uniformly fatal lung disease marked by destruction and scarring of the lung parenchyma and progressive loss of respiratory function. IPF affects nearly 3 million people worldwide, and annual mortality in the US alone exceeds 40,000. Nintedanib and pirfenidone, the only drugs approved for the treatment of IPF, slow progression but do not cure the disease. Consequently, there is a pressing need for effective treatments beside lung transplantation. Unfortunately, predictive models of IPF are not available, underscoring the critical need for physiologically relevant *in-vitro* substrates that enable quantitative and mechanistic studies of human IPF. Here we report the development and characterization of a human pulmonary fibrosis-specific cell culture substrate comprised of intact fibrotic lung extracellular matrix that recapitulates the human IPF disease environment *in vitro*. We document the activation and disease-specific phenotype of human lung fibroblasts cultured in the IPF disease-specific substrate, and establish feasibility of testing antifibrotic agents using this substrate. Altogether, our results demonstrate the applicability of this fibrosis-specific substrate for 3D *in-vitro* models of IPF and cell-based assays in early-stage drug discovery.

## INTRODUCTION

Idiopathic pulmonary fibrosis (IPF) is a chronic interstitial lung disease that primarily affects older adults and is associated with dysregulation of pulmonary fibroblasts, extensive remodeling and deposition of extracellular matrix, and progressive loss of respiratory function.^1–3^ Incidence and prevalence appear to be increasing worldwide with aging populations and improved diagnostics.^4^ Every year more than 50,000 new patients are diagnosed with IPF^5^ – an incidence comparable to those of liver, stomach, testicular, or cervical cancers.^6^ After diagnosis, median survival is only 3 – 4 years, and annual mortality exceeds 40,000.^7^ The etiology of IPF remains unknown, but risk factors include smoking, environmental exposures, chronic viral infections, gastroesophageal reflux, lung injury, and genetic predispositions.^4, 8^ Nintedanib and pirfenidone, the only drugs approved to treat IPF, attenuate disease progression but do not prevent decline,^1, 4, 9^ necessitating the development of new drugs that can effectively treat IPF.

A major obstacle to developing effective treatments for IPF is the lack of predictive animal and *in-vitro* models of IPF. Animal models of pulmonary fibrosis are well-established in rodents^10–12^ but present fibrosis that resolves over time rather than the progressive, non-resolving fibrotic process characteristic of IPF in humans.^3, 13^ Furthermore, there are no robust or widely adopted *in-vitro* models of IPF to enable predictive basic and translational studies.^14^ Consequently, an *in-vitro* model of IPF that emulates human pathophysiology could enable critical new insights into the natural history and pathological mechanisms of IPF, and guide therapeutic development.

Current *in-vitro* models of IPF have limited physiologic relevance because they fail to recapitulate the complex biochemical, structural, and mechanical environment of fibrotic human lungs. In fibrosis, the extracellular matrix (ECM) has different biochemical composition, stores more fibrogenic growth factors, and has altered structure and biomechanics compared to normal ECM,^15–17^ and the direct influence of growth factors^18, 19^ and increased matrix stiffness^20^ on myofibroblast differentiation has been previously demonstrated. Altogether, such matrix alterations induce a profibrotic microenvironment, activate pulmonary fibroblasts, and suggest that IPF progression is correlated with an abnormal ECM microenvironment.^21^ As lung matrix is implicated in both lung function and fibrotic disease progression, IPF models and drug screening platforms not incorporating lung ECM lack defining components of the IPF disease environment. The most common *in-vitro* IPF drug testing platforms utilize cell culture plates coated with collagen type I and culture media supplemented with high concentrations of transforming growth factor β (a profibrotic cytokine associated with fibrogenesis),^22^ but no testing platforms that utilize other IPF disease-specific ECM components have been established.

An *in-vitro* cell culture substrate comprised of fibrotic human lung matrix could faithfully recapitulate the composition, structure, and mechanics of the human IPF disease environment. While removal of native cells (decellularization) from human tissues has been demonstrated in a number of tissues including lungs^23–26^, efforts have been primarily focused on the isolation and characterization of ECM from normal, non-diseased tissues. Reproducible, scalable methods for the production of disease-specific ECM biomaterials from diseased human tissues such as fibrotic lungs have not been robustly established. Furthermore, an *in-vitro* cell culture substrate that recapitulates the complex disease environment of human IPF tissue would be an extremely valuable tool for screening antifibrotic agents in early-stage development.

In this study, we investigated the feasibility of developing a cell culture substrate from fibrotic human lung tissue for 3D *in-vitro* models of human pulmonary fibrosis. Our hypothesis was that normal human lung fibroblasts would display a disease-specific phenotype *in vitro* in the presence of fibrotic lung extracellular matrix. We implemented a ‘physiomimetic approach’ to develop disease-specific IPF cell culture substrates (scaffolds) comprised of lung extracellular matrix derived from human IPF tissues (Fig. 1). Our goal was to develop a human fibrotic lung ECM biomaterial for use as a 3D cell culture substrate for predictive *in-vitro* models of IPF that could reduce dependence on animal models while enabling physiologically relevant results. Such disease-specific cell culture substrates could radically improve the physiological relevance of *in-vitro* models of IPF and antifibrotic drug screening platforms, and accelerate development of safe and effective IPF treatments.

**Figure 1.**
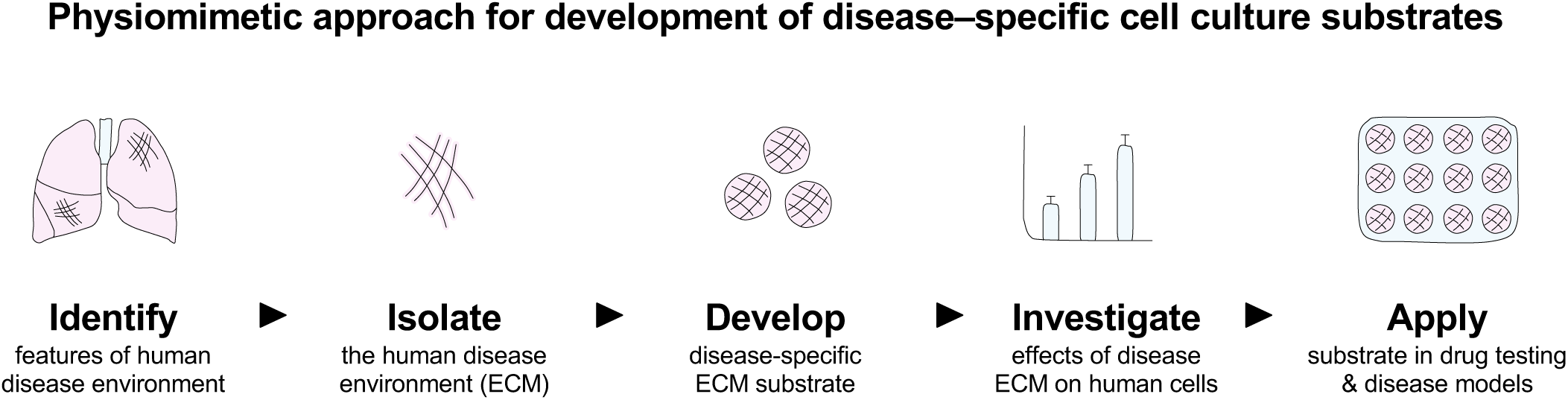
Overview of physiomimetic approach. Our development of IPF disease-specific cell culture substrates is guided by a physiomimetic approach that aims to identify and isolate the human disease environment, then develop and investigate disease-specific ECM substrates *in vitro* utilizing disease-relevant human cell types (e.g., pulmonary fibroblasts) whose phenotype can be directly compared against diseased human IPF lung specimens prior to application in IPF disease models and antifibrotic drug testing.

## MATERIALS & METHODS

### Procurement of human lung tissues

Acceptance criteria for donors of normal and IPF lungs were established prior to initiation of studies. Normal lung donors had no history, diagnosis, or evidence of: smoking, aspiration pneumonia, asthma, chronic obstructive pulmonary disease, cystic fibrosis, emphysema, interstitial lung disease, or lung cancer. IPF donors required diagnosis of idiopathic pulmonary fibrosis confirmed by a lung transplant pathologist. All IPF donors had end-stage disease and were recipients of lung transplants. Normal human lungs (*n* = 3) not acceptable for use in transplantation were procured under a protocol approved by the Institutional Review Board at the International Institute for the Advancement of Medicine. Diseased human lungs (*n* = 3) designated as surgical waste were procured under protocols approved by the Institutional Review Boards at Vanderbilt University Medical Center and State University of New York (SUNY) Downstate Medical Center. Lungs were procured in standard fashion, flushed with cold organ preservation solution, transported on ice, and made available without identifiers. In this study, to minimize variability, lung tissues from right middle and right lower lobes were utilized.

### Characterization of lung donors

Lung donor characteristics were tabulated from deidentified summaries provided by the United Network of Organ Sharing (UNOS) under approved protocols and in compliance with all applicable regulations.

### Sampling of lung tissues

Tissue samples were collected from medial, lateral, and peripheral regions of right middle lobes (2 samples per region), for a total of 6 regional samples per right middle lobe, and 18 regional samples each for IPF and normal lungs.

### Preparation of lung matrix scaffolds

Upon receipt, lungs were rinsed with cold sterile saline. Native lung tissue samples were collected for histologic analyses, then lung tissues were stored at −80°C. At the time of use, lung tissues were processed under sterile conditions with a proprietary combination of chemicals, enzymes, and surfactants to remove cellular components and isolate normal and fibrotic lung extracellular matrix. Matrix scaffolds (diameter: 7 mm, thickness: 1 mm) were prepared under sterile conditions for experimental use. For all assays, three tissue samples or matrix scaffolds were randomly selected from each lung and evaluated in triplicate.

### Histologic analyses of lung tissues and scaffolds

Lung tissue samples were fixed in cold phosphate-buffered 4% paraformaldehyde for 24 hours, embedded in paraffin, and sectioned at 5 or 10 μm thickness. Three sections (medial, lateral, peripheral) from all normal and fibrotic lungs were stained with hematoxylin and eosin, trichrome, Verhoeff–Van Gieson, Alcian blue, and pentachrome, and examined under light microscopy. Representative images were obtained using a fluorescence microscope (FSX100, Olympus).

### Histopathologic characterization of lung tissues

All lung sections were subjected to blinded review by a lung transplant pathologist. Slides were randomized, arbitrarily numbered, and delivered without reference to the pathologist, who reviewed and assigned fibrosis scores to all regions in 5 high-power fields according to a standard pulmonary fibrosis scoring rubric^27^ to quantify the extent of architectural disruption and fibrosis (Supplementary Fig. 1C). Fibrosis scores from each high-power field were averaged to obtain an average fibrosis score for each region of lung. To quantitatively assess the severity and distribution of fibrosis, a grid with unit length 250 µm was overlaid onto each high-power field (20×) image, and regions corresponding to various classifications of fibrosis were outlined (Supplementary Fig. 1F). Each region was assigned a calculated relative percent area of the high-power field using the grid. Each fibrosis score was weighted according to percent area, and average fibrosis scores for each high-power field were calculated based on the weighted average of all regional fibrosis scores in each high-power field. Fibrosis scores were then averaged across 5 high-power fields per region, with four regions evaluated per lobe. Only regions of IPF lungs with confirmed fibrosis score ≥ 2 were investigated in this study (Supplementary Fig. 1D,E,G).

### Biochemical characterization of lung tissues and scaffolds

To quantify collagen in lung tissues and scaffolds, samples were weighed, homogenized, and digested with pepsin (0.1 mg mL^-1^) in 0.5M acetic acid for 12 hours at 4°C, and subjected to a collagen quantification assay (Sircol, Biocolor) according to the manufacturer’s instructions. To quantify sulfated glycosaminoglycans in lung tissues and scaffolds, samples were weighed, homogenized, and digested with papain (1 μg mL^-1^) for 12 hours at 60°C, and subjected to the dimethylene blue dye assay, wherein absorbance was measured at 595 nm. To quantify elastin in lung tissues and scaffolds, samples were weighed and homogenized, and soluble α–elastin was extracted via three extractions with hot 0.25M oxalic acid. Samples were then subjected to an elastin quantification assay (Fastin, Biocolor) according to the manufacturer’s instructions. To quantify residual DNA in matrix scaffolds, samples were subjected to a quantitative DNA assay (Quant-iT PicoGreen, Invitrogen) according to the manufacturer’s instructions.

### Immunohistochemical staining

Following de-paraffinization, sections of lung tissues and scaffolds were subjected to boiling citrate buffer (pH 6.0) for antigen retrieval, and blocked with 5% normal goat serum in phosphate-buffered saline for 1 hour at room temperature. Next, antibodies were diluted as necessary, applied, and incubated for 12 hours at 4°C or 4 hours at room temperature. Sections were mounted (VectaMount Permanent Mounting Medium, Vector Laboratories), and coverslips were applied. Images were obtained using a light microscope (Eclipse Ts2, Nikon). Immunohistochemical stains were performed for alpha smooth muscle actin (Cell Signaling Technology, 19245), fibrillin 2 (Sigma Life Science, HPA012853), Ki67 (ThermoFisher Scientific, PA1-38032), laminin γ1 (Abcam, ab233389), matrix gla protein (LS Bio, LS-B14824), and periostin (Abcam, ab14041). A list of antibodies with dilutions used is provided in Supplementary Table 1.

**Table 1.**
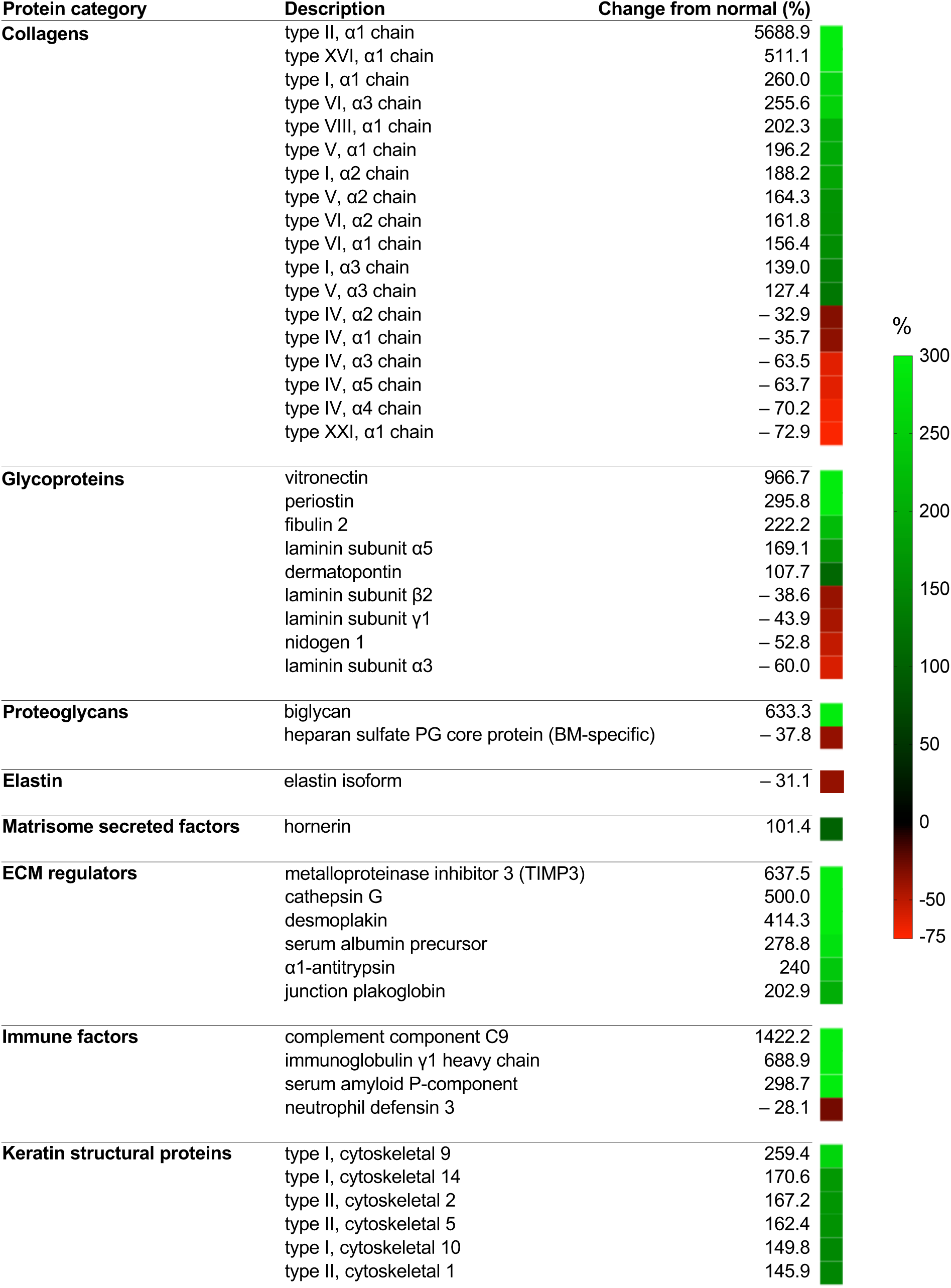
Mass spectrometry analysis of IPF lung matrisome. Changes from normal in the abundance of IPF lung matrisome components. PG: proteoglycan, BM: basement membrane.

| Protein category | Description | Change from normal (%) |  |
| --- | --- | --- | --- |
| **Collagens** | type II, α1 chain | 5688.9 | |
| | type XVI, α1 chain | 511.1 | |
| | type I, α1 chain | 260.0 | |
| | type VI, α3 chain | 255.6 | |
| | type VIII, α1 chain | 202.3 | |
| | type V, α1 chain | 196.2 | |
| | type I, α2 chain | 188.2 | |
| | type V, α2 chain | 164.3 | |
| | type VI, α2 chain | 161.8 | |
| | type VI, α1 chain | 156.4 | |
| | type I, α3 chain | 139.0 | |
| | type V, α3 chain | 127.4 | |
| | type IV, α2 chain | -32.9 | |
| | type IV, α1 chain | -35.7 | |
| | type IV, α3 chain | -63.5 | |
| | type IV, α5 chain | -63.7 | |
| | type IV, α4 chain | -70.2 | |
| | type XXI, α1 chain | -72.9 | |
| **Glycoproteins** | vitronectin | 966.7 |  |
|  | periostin | 295.8 |  |
|  | fibulin 2 | 222.2 |  |
| | laminin subunit α5 | 169.1 | |
|  | dermatopontin | 107.7 |  |
| | laminin subunit β2 | -38.6 | |
| | laminin subunit γ1 | -43.9 | |
|  | nidogen 1 | -52.8 |  |
| | laminin subunit α3 | -60.0 | |
| **Proteoglycans** | biglycan | 633.3 |  |
|  | heparan sulfate PG core protein (BM-specific) | -37.8 |  |
| **Elastin** | elastin isoform | -31.1 |  |
| **Matrisome secreted factors** | hornerin | 101.4 |  |
| **ECM regulators** | metalloproteinase inhibitor 3 (TIMP3) | 637.5 |  |
|  | cathepsin G | 500.0 |  |
|  | desmoplakin | 414.3 |  |
|  | serum albumin precursor | 278.8 |  |
| | α1-antitrypsin | 240 | |
|  | junction plakoglobin | 202.9 |  |
| **Immune factors** | complement component C9 | 1422.2 |  |
| | immunoglobulin γ1 heavy chain | 688.9 | |
|  | serum amyloid P-component | 298.7 |  |
|  | neutrophil defensin 3 | -28.1 |  |
| **Keratin structural proteins** | type I, cytoskeletal 9 | 259.4 |  |
|  | type I, cytoskeletal 14 | 170.6 |  |
|  | type II, cytoskeletal 2 | 167.2 |  |
|  | type II, cytoskeletal 5 | 162.4 |  |
|  | type I, cytoskeletal 10 | 149.8 |  |
|  | type II, cytoskeletal 1 | 145.9 |  |

### Quantification of immunohistochemical staining by image analysis

Images of immuno-histochemical stains were captured using a slide scanner (P250 High Capacity Slide Scanner, 3D Histech). To quantify immunohistochemical staining, images were analyzed using an image analysis software module (DensitoQuant, Quant Center, 3D Histech), and the number of positive and negative pixels were quantified and analyzed.

### Mass spectrometry of IPF and normal lung matrisomes

Detailed methods are available in the Supplementary Information.

### Quantification of growth factors

To quantify growth factors in native lung tissues and matrix scaffolds, a multiplex growth factor array (Quantibody Human Growth Factor Array Q1; Ray Biotech) was performed and analyzed by Q-Analyzer software. To quantify growth factors secreted by human fibroblasts *in vitro*, enzyme-linked immunosorbent assays (ELISA) were performed for bFGF (R&D Systems, DFB50) and TGFβ (R&D Systems, DB100B). All samples were analyzed in triplicate.

### Scanning electron microscopy

Lung matrix samples were collected, fixed in formalin for 24 hours, rinsed in 70% ethanol, frozen, lyophilized, and imaged using an electron microscope (GeminiSEM 300, Zeiss) with accelerating voltage 2.5 kV.

### Transmission electron microscopy

Lung matrix samples were fixed with 2.5% glutaraldehyde, 4% paraformaldehyde, and 0.02% picric acid in 0.1M Na-cacodylate buffer (pH 7.2). Samples were then post-fixed with 1% OsO4 in Sorenson’s buffer for 1 hour, dehydrated, and embedded in Lx-112 (Ladd Research Industries). Sections (thickness: 60 nm) were prepared using a PT-XL ultramicrotome, stained with uranyl acetate and lead citrate, and examined with an electron microscope (JEM-1200 EXII; JEOL). Images were captured with a digital camera (ORCA-HR; Hamamatsu Photonics) and recorded with imaging software (Image Capture Engine, AMT).

### Mechanical testing of IPF and normal lung scaffolds

Uniaxial tensile mechanical testing was conducted with a 10 N load cell (Model 5848, Instron), as previously described.^23^ Lung tissues and matrix from transverse sections of the right middle lobe were randomly selected and dissected into 3 cm by 1 cm samples. A consistent orientation from right middle lobe was maintained to minimize effects of lung anisotropy on mechanical testing data. Samples were secured and mounted, and a pre-load of 0.003 N was applied. All samples were tested at the same grip-to-grip distance for consistency. Samples were kept hydrated throughout all mechanical testing with phosphate-buffered saline at room temperature. A 20% uniaxial strain was applied at a strain rate of 1% s^-1^, and at frequencies of 0.25, 0.50, or 0.75 Hz.

### Cell culture

Human lung fibroblasts (ATCC) were cultured in Dulbecco’s Modified Eagle Medium (DMEM) supplemented with 10% fetal bovine serum and 1% penicillin/streptomycin under standard culture conditions with 5% CO_2_ at 37°C.

### Gene expression analysis

Total RNA was extracted (RNeasy Micro Kit, QIAGEN), and cDNA synthesis was performed using random primers (iScript Select cDNA Synthesis Kit, Bio-Rad). Quantitative real-time polymerase chain reaction (qPCR) was performed in triplicate using master mix (Brilliant III Ultra-Fast SYBR Green QPCR Master Mix, Agilent Technologies) and a real-time PCR system (AriaMax Real PCR System, Agilent Technologies). A list of primers is provided in Supplementary Table 2.

### Drug testing

Normal human fibroblasts were cultured *in vitro* for 24 hours, then exposed to antifibrotic agent PF3644022 hydrate (PZ-0188, Sigma-Aldrich) at a concentration of 1 µM for 72 hours. Metabolic activity was measured using Alamar Blue reagent (DAL1025, ThermoFisher Scientific) according to the manufacturer’s instructions. The reagent was added to cells in culture at 24, 48 and 72 hours, and incubated for 4 hours before readout. Absorbance was measured at 570nm, with reference wavelength at 600 nm.

### Statistical analyses

One-way ANOVA and Student’s *t*-tests were performed using statistical analysis software (Prism 8, GraphPad), and *p* < 0.05 was considered significant.

## RESULTS

### Assessment of IPF and normal lungs

Donor characteristics of IPF and normal lung tissues were analyzed to confirm that there were no significant differences in age, height, weight, body mass index, or smoking history (Supplementary Fig. 1A,B; Supplementary Table 3). An established numerical rubric^27^ was used to assess the extent of histomorphologic disruption and fibrosis. Tissue sampling and histopathologic analyses are described in detail in Supplementary Information.

### Preparation of lung matrix scaffolds

Native lung tissues were treated with a proprietary combination of chemicals, enzymes, and surfactants to remove cellular and nuclear components, which was confirmed by hematoxylin and eosin staining (Fig. 2A) and quantitative DNA assay (Supplementary Fig. 2A). Matrix scaffolds from all human lungs were confirmed negative for mycoplasma, bacteria, and fungi (Supplementary Fig. 2B), and deemed suitable for use in cell-based studies.

**Figure 2.**
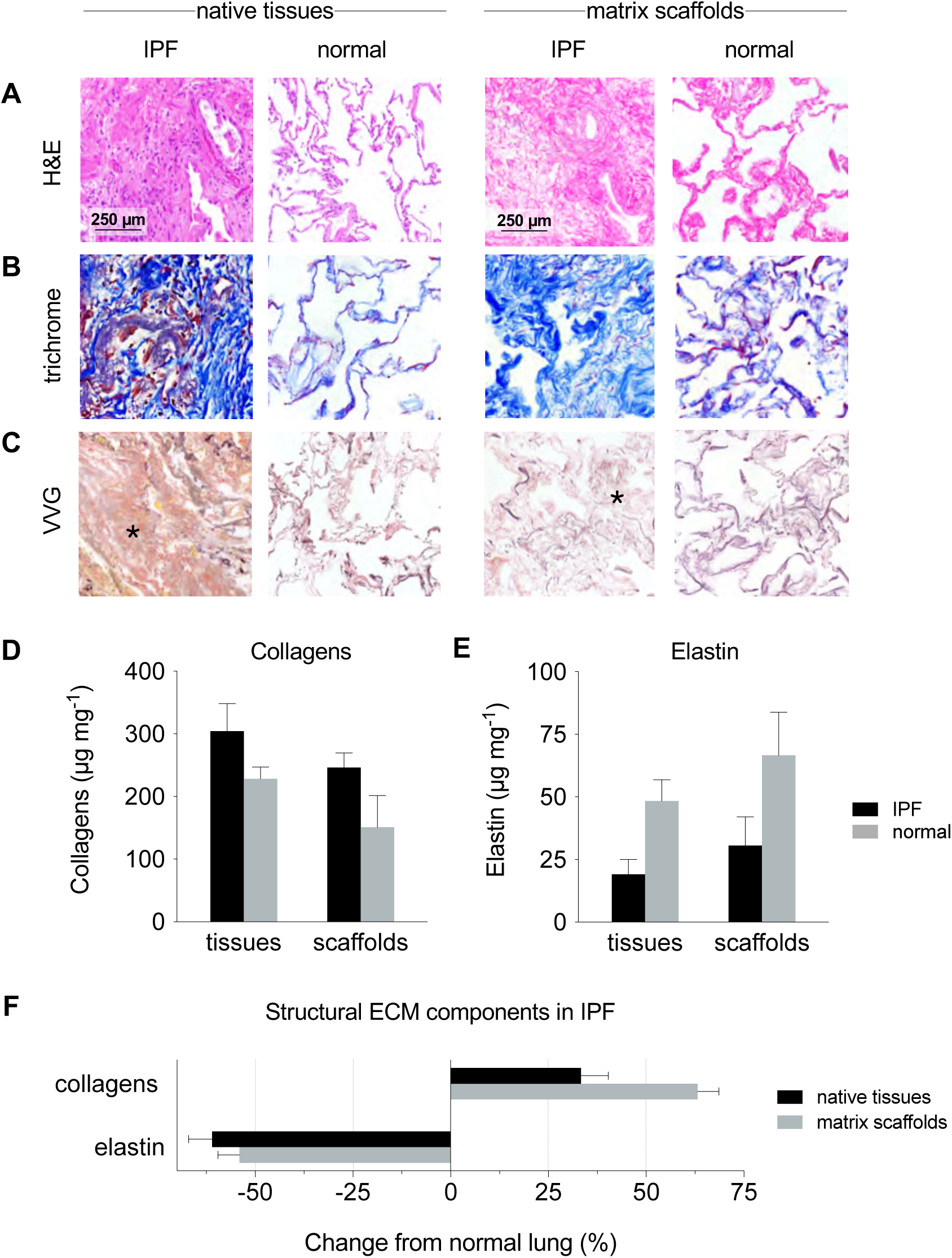
Histological & biochemical characterization of ECM structural components in IPF lung tissues and matrix scaffolds. Representative micrographs of histologic stains: (**A**) hematoxylin and eosin (H&E), (**B**) trichrome (collagens, blue), and (**C**) Verhoeff–Van Gieson (VVG, elastic fibers, black) demonstrating differences in histomorphology of IPF and normal lung tissues and scaffolds. Star indicates representative region with severe fibrosis and loss of elastic fibers. Quantification of structural ECM components (**D**) collagens and (**E**) elastin by biochemical assays. (**F**) Changes from normal lung in structural ECM components in IPF. H&E: hematoxylin and eosin, VVG: Verhoeff–Van Gieson.

### IPF matrix scaffolds recapitulate disease-specific histologic features

For histologic evaluations of IPF, representative fields corresponding to fibrosis score 3 (severe fibrosis) were selected. To visualize distributions of ECM structural components in IPF and normal lungs, histologic staining was performed on native (untreated) tissues and matrix scaffolds. H&E staining of native IPF tissues revealed severe distortion of lung structure and large areas of fibrous obliteration with minimal remaining airspace (Fig. 2A). By contrast, H&E staining of native normal lung tissues displayed abundant airspaces defined by thin alveolar septa and stereotypical alveolar saccular architecture. Matrix scaffolds from analogous regions of IPF and normal lungs had no discernible nuclei and displayed drastic differences in scaffold architecture consistent with fibrotic and normal native lung tissues, respectively. Trichrome staining showed dramatic deposition of collagens (blue) throughout regions of severe fibrosis (Fig. 2B). In IPF tissues and scaffolds, collagen fibers were observed in densely aligned bundles and in loosely disorganized networks; whereas in normal lung tissues and scaffolds, collagen was organized along alveolar septa and within the interstitium. Verhoeff–Van Gieson (VVG) elastic staining showed a notable loss of elastic fibers (black) in regions of IPF tissues and scaffolds with severe fibrosis, whereas in normal lung tissues and scaffolds elastic fibers were dispersed homogenously throughout the respiratory zone (Fig. 2C).

### IPF matrix scaffolds contain disease-specific biochemical composition

Soluble collagens were quantified in native tissues and matrix scaffolds, and increases in collagens were measured relative to normal in IPF native tissues (33.3 ± 19.2 %) and matrix scaffolds (63.2 ± 15.6 %, Fig. 2D). Consistent with the loss of elastic fibers observed in VVG elastic staining, quantification of elastin confirmed reduction in IPF native tissues (60.6 ± 12.3 %) and matrix scaffolds (54.1 ± 17.2 %) relative to normal (Fig. 2E). Altogether, the structural ECM components in IPF demonstrated clear trends relative to normal in both native tissues and matrix scaffolds: increased collagens (33 – 63%) and decreased elastin (54 – 61%; Fig. 2F). Alcian blue and pentachrome staining were performed to assess the extent and distribution of proteoglycans in IPF tissues, which was significantly higher in areas of moderate and severe fibrosis (scores ≥ 2) than in areas of mild fibrosis (scores < 2) and normal lung tissues (Fig. 3A,B). Quantification of sulfated glycosaminoglycans (GAG) revealed that GAG components in IPF native tissues and scaffolds was 232.5 – 300.5% higher than in normal lungs (Fig. 3C-E), consistent with overexpression of sulfated glycosaminoglycans previously observed in fibrotic foci^28^. Immunohistochemical staining of IPF tissues for multiple ECM glycoproteins revealed dramatic differences from normal lung tissues in fibrillin 2, laminin γ1, matrix GLA protein (MGP), and periostin (Fig. 3F-I). Areas with severe fibrosis (fibrosis score: 3) were characterized by pervasive overexpression of fibrillin 2, MGP, and periostin, and loss of laminin γ1. Notably, changes from normal lung were consistent in native tissues and matrix scaffolds for all glycoproteins that were investigated (Fig. 3J).

**Figure 3.**
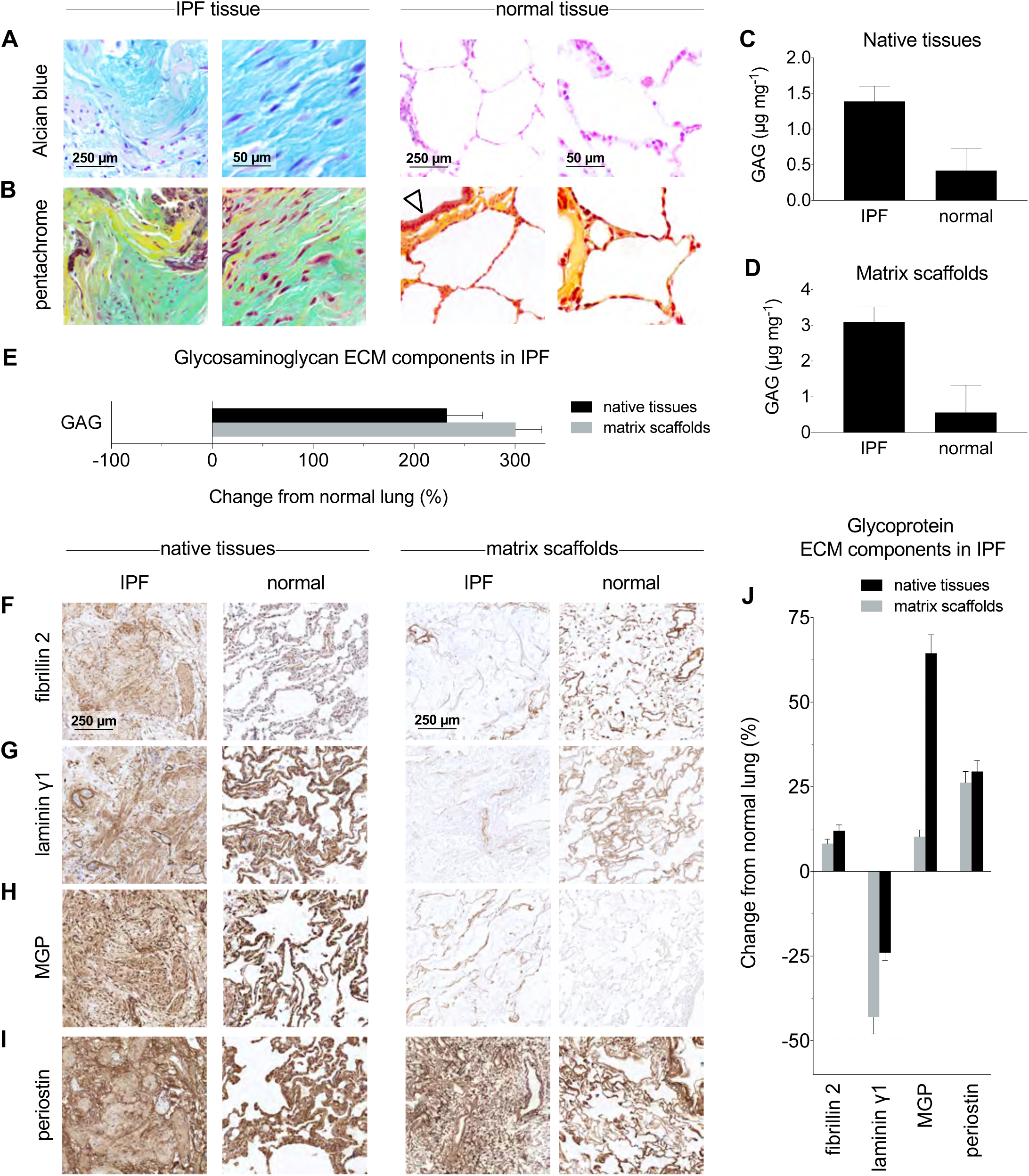
Characterization of proteoglycans and glycoproteins in IPF lung tissues and matrix scaffolds. Representative micrographs of histologic stains: (**A**) Alcian blue (proteoglycans, blue) and (**B**) pentachrome (acidic polysaccharides, green) demonstrating differences in proteoglycans between IPF and normal lung tissues. Arrow indicates normal airway epithelium. Quantification of sulfated glycosaminoglycan ECM components in (**C**) native tissues and (**D**) matrix scaffolds. (**E**) Changes from normal lung in glycosaminoglycan ECM components in IPF. Immunohistochemical staining of glycoprotein ECM components in IPF: (**F**) fibrillin 2, (**G**) laminin γ1, (**H**) matrix gla protein (MGP), (**I**) periostin. (**J**) Quantification of glycoproteins by image analysis of immunohistochemical staining using DensitoQuant software.

Mass spectrometry was performed on IPF and normal lung matrix scaffolds to assess the IPF matrisome (Table 1), and revealed changes from normal lung consistent with histopathologic observations and biochemical assays. Multiple collagen types increased above 150%, including collagen types I, II, V, VI, VIII, XVI. Notably, in IPF lungs collagen types IV and XXI – the primary collagens comprising the alveolar basement membrane – decreased between 33 – 73%, consistent with the loss of basement membrane and alveolar structure associated with the progression of pulmonary fibrosis.^29^ The glycoprotein vitronectin was elevated 967%, and glycoproteins fibulin 2 and periostin were both elevated above 200%. Laminin subunits α3, β2, γ1, and nidogen 1, which are associated with the basement membrane, were all decreased in IPF lungs. Biglycan was increased by 633%, however basement membrane-specific heparan sulfate proteoglycan core protein was decreased by 38%. Elastin isoforms were also decreased by 31%, consistent with quantitative biochemical analyses. Interestingly, in IPF lungs several regulators of the extracellular matrix were also increased more than 200% above normal, including metalloproteinase inhibitor 3 (TIMP3), cathepsin G, desmoplakin, and α1-antitrypsin.

To assess changes in endogenous growth factors, a multiplex growth factor array was performed. Two growth factors were detected only in IPF native tissues and not in normal lung native tissues: transforming growth factor beta 3 (TGF-β3) and heparin-binding EGF-like growth factor (HB-EGF; Supplementary Table 4). In IPF native tissues, insulin-like growth factor binding protein 1 (IGFBP-1) was 160-fold above normal, and both basic fibroblast growth factor (bFGF) and endocrine gland-derived vascular endothelial growth factor (EG-VEGF) were approximately 20-fold above normal. Brain-derived neurotrophic factor (BDNF) and growth differentiation factor 15 (GDF-15, a prognostic factor for IPF^30^) were elevated 3-fold to 5-fold, but osteoprotegerin (OPG) was reduced by more than half. Five growth factors were detected in IPF matrix scaffolds (Table 2), including IGFBP-6, whose family of carrier proteins were shown to induce production of collagen type I and fibronectin in normal primary lung fibroblasts^31, 32^. Neurotrophin-4 (NT-4), which is elevated in explanted IPF lungs and shown to drive proliferation of primary human lung fibroblasts through TrkB-dependent and protein kinase B-dependent pathways^33^, was also detected in IPF matrix scaffolds.

**Table 2.** Quantification of growth factors in IPF and normal lung matrix scaffolds. Growth factor concentrations were measured by multiplex growth factor array. Green arrow (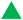) indicates positive fold change (increase) from normal in concentration of growth factors. Red arrow (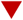) indicates negative fold change (decrease) from normal in concentration of growth factors. ND: not detected.

| Growth factor | Description | Concentration (pg mL<sup>-1</sup>) |  | Fold change from normal |
| --- | --- | --- | --- | --- |
|  |  | Normal scaffold | IPF scaffold |  |
| GDF-15 | Growth differentiation factor 15 | 0.8 | 14.5 | + 18.1 ▲ |
| BDNF | Brain-derived neurotrophic factor | 6.3 | 48.7 | + 7.7 ▲ |
| IGFBP-6 | Insulin-like growth factor binding protein 6 | 14.1 | 71.8 | + 5.1 ▲ |
| HGF | Hepatocyte growth factor | 23.8 | 91.0 | + 3.8 ▲ |
| EG-VEGF | Endocrine gland-derived vascular endothelial growth factor | 0.6 | 1.5 | + 2.5 ▲ |
| bFGF | Basic fibroblast growth factor | 22.6 | 54.8 | + 2.4 ▲ |
| HB-EGF | Heparin-binding EGF-like growth factor | 1.4 | 2.5 | + 1.8 ▲ |
| TGF-β3 | Transforming growth factor β3 | 2.8 | 4.0 | + 1.4 ▲ |
| VEGF | Vascular endothelial growth factor | 4.6 | 3.0 | − 0.3 ▼ |
| EGF R | Epidermal growth factor receptor | ND | ND | – |

### IPF matrix scaffolds have disease-specific structural and mechanical properties

The gross appearance of IPF matrix scaffolds was dramatically different from the appearance of normal lung matrix scaffolds. Normal lung matrix scaffolds appeared translucent, with visible bronchial and vascular conduits and saccular structures throughout the parenchyma (Fig. 4A). By contrast, IPF matrix scaffolds had pervasive dense fibroconnective structures, with abnormal disorganized architecture, honeycombing, and no apparent airways or vessels. Scanning electron microscopy revealed dramatic disruption of normal alveolar architecture in IPF scaffolds (Fig. 4B). Topography of collagen fibers in IPF scaffolds was visualized by inverted color micrographs of trichrome staining, which showed dense fibrous bundles in IPF scaffolds and stereotypical porous (alveolar-like) networks in normal lung scaffolds (Fig. 4C). Transmission electron microscopy showed dense fibrous bands (F) of extracellular matrix in IPF matrix scaffolds with minimal evidence of normal basement membrane, whereas normal lung matrix scaffolds had abundant airspaces (A), delicate basement membrane (arrow), and alveolar capillaries (C; Fig. 4D). Uniaxial mechanical testing of IPF and normal tissues and scaffolds indicated that IPF tissues and scaffolds were approximately 20× stiffer at 5% strain and approximately 5× stiffer at 20% strain compared to normal tissues and scaffolds (Fig. 4E). Importantly, mechanical testing also confirmed that the processing of native tissues to obtain matrix scaffolds did not alter the mechanical properties of matrix scaffolds from native tissues, as differences in elastic modulus between native tissues and matrix scaffolds were not significant (Fig. 4F,G).

**Figure 4.**
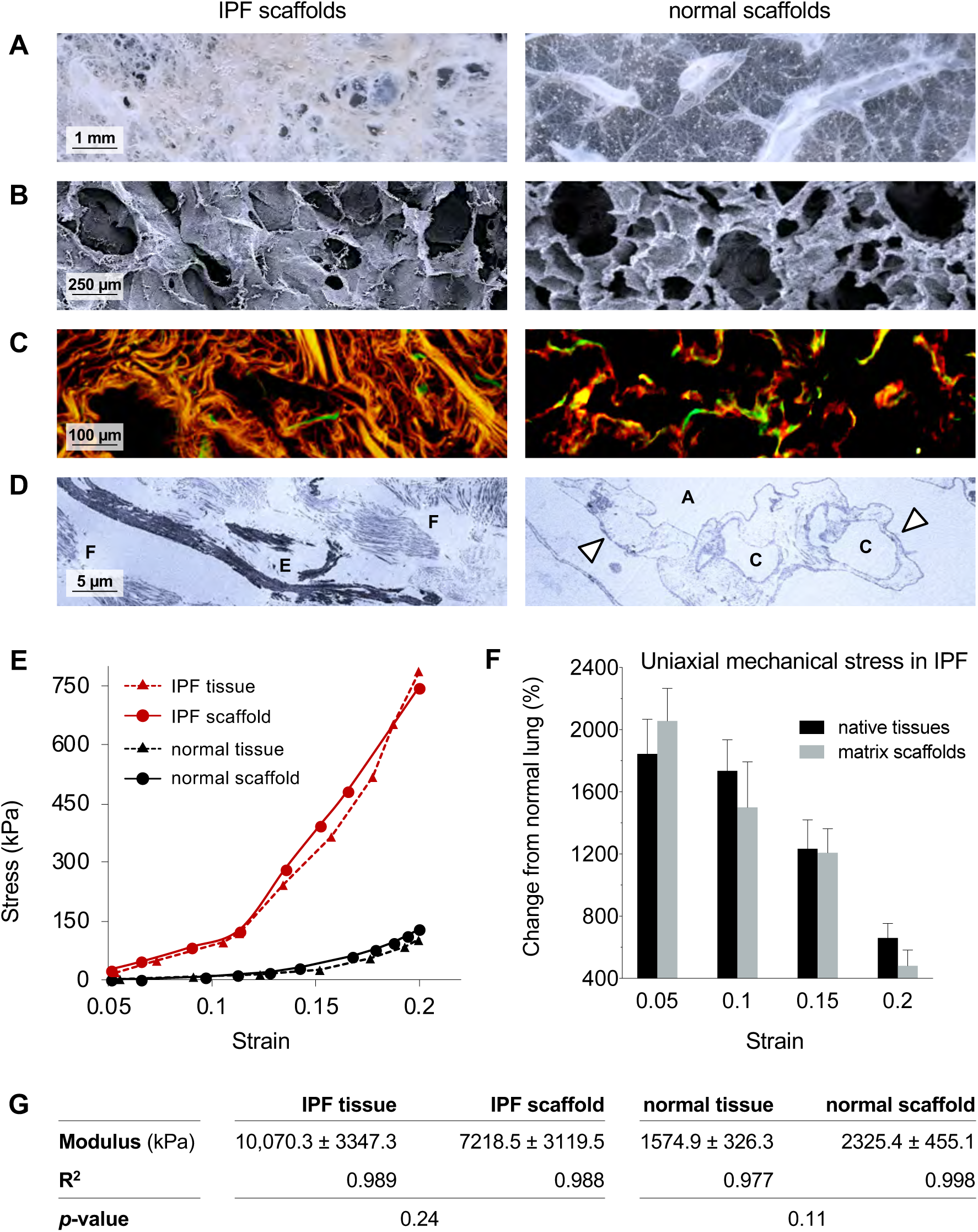
Structural, topographical, and mechanical characterization of IPF lung scaffolds. Representative images of IPF and normal lung scaffolds: (**A**) gross photography, (**B**) scanning electron microscopy, (**C**) light microscopy (inverted color micrograph) of trichome staining demonstrating topography of ECM fibers in IPF scaffolds, (**D**) transmission electron microscopy. A: airspace, C: alveolar capillary, E: elastin bundle fragments, F: fibroconnective collagenous matrix, arrow: basement membrane. (**E**) Representative uniaxial stress-strain curves of IPF and normal lung tissues and matrix scaffolds. (**F**) Change in uniaxial mechanical stress from normal lung tissues and matrix scaffolds. (**G**) Tangent modulus values. Statistical analyses between tissues and scaffolds were performed using Student’s *t*-test, with significance when *p* < 0.05. All values represent mean ± standard deviation.

### IPF matrix scaffolds support disease-like phenotype of lung fibroblasts

Normal human lung fibroblasts were added to IPF and normal lung matrix scaffolds and cultured *in vitro* for 7 days. H&E staining showed that the phenotype of normal human lung fibroblasts varied between cells cultured in IPF and normal lung matrix scaffolds (Fig. 5A). Fibroblasts in IPF matrix scaffolds showed higher expression of alpha smooth muscle actin than fibroblasts in normal lung matrix scaffolds. Morphologic similarities between fibroblasts cultured in IPF scaffolds and IPF native tissue were observed (Fig. 5B). In contrast, immunostaining of FOXO3, a transcription factor whose downregulation is linked to fibrogenesis^34^, showed lower expression in human lung fibroblasts cultured on IPF matrix scaffolds compared to fibroblasts cultured on normal lung matrix scaffolds (Fig. 5C). Consistent with alpha smooth muscle immunohistochemical staining, gene expression analysis showed significant upregulation of ACTA2 (alpha smooth muscle actin). Additional upregulated fibrosis-specific markers of fibroblast activation included COL1A1 (collagen type I, subunit α1), MMP2, PDGFC, PTEN, and PRRX1 (Fig. 5D). Activation of fibroblasts *in vitro* was also assessed by quantification of secreted basic fibroblast growth factor (bFGF) and transforming growth factor beta (TGFβ), with normal human lung fibroblasts cultured on tissue culture plastic as a standard control. Interestingly, secretion of bFGF and TGFβ were both highest with fibroblasts cultured in IPF matrix scaffolds (Fig. 5E,F). Notably, secreted TGFβ was significantly higher in IPF matrix scaffolds compared to normal lung matrix scaffolds, suggesting that substrate stiffness may have influenced secretion of TGFβ.

**Figure 5.**
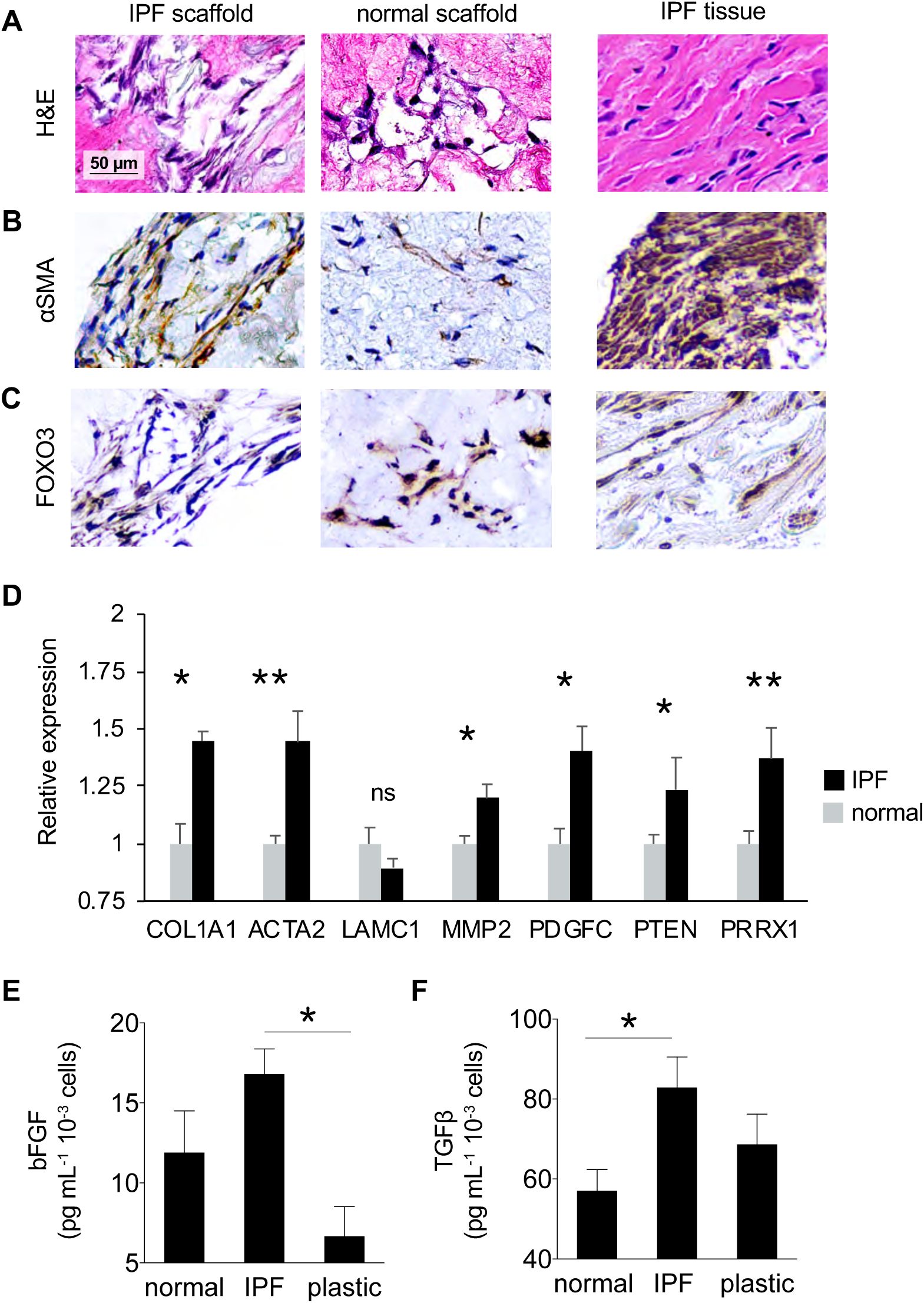
Phenotype of lung fibroblasts in IPF and normal lung scaffolds. Representative micrographs of (**A**) H&E and immunohistochemical staining of (**B**) alpha smooth muscle actin (αSMA) and (**C**) Forkhead box O3 (FOXO3). (**D**) Gene expression of normal human lung fibroblasts cultured in IPF and normal lung scaffolds. * *p* < 0.05, ** *p* < 0.01, ns: not significant. Quantification by ELISA of (**E**) basic fibroblast growth factor (bFGF, * *p* < 0.05) and (**F**) transforming growth factor beta (TGFβ, * *p* < 0.05) secreted by normal human lung fibroblast cultured in IPF and normal lung scaffolds and on tissue culture plastic. All values represent mean ± standard deviation.

### IPF matrix scaffolds provide a disease-specific environment for testing antifibrotic agents

Pulmonary fibroblasts in IPF matrix scaffolds showed a mean growth rate (linear fit: slope = 6.74, R^2^ = 0.98) over 80% faster than fibroblasts in normal lung matrix scaffolds (linear fit: slope = 3.70, R^2^ = 0.93; Fig. 6A, no drug), consistent with the fibroproliferative process characteristic of human IPF. To assess differences in phenotype between fibroblasts cultured on IPF matrix scaffolds and the conventional drug testing substrate tissue culture plastic, disease-associated gene expression and growth factor secretion were analyzed. Fibroblasts cultured in IPF matrix scaffolds expressed significantly higher COL1A1 and MMP2 than fibroblasts cultured on plastic (Fig. 6B), and secreted more profibrotic growth factors bFGF and TGFβ than fibroblasts cultured in normal lung matrix or on plastic (Fig. 6C,D), suggesting that the presence of disease-specific matrix resulted in more disease-associated fibroblast phenotype *in vitro* compared to fibroblasts on plastic. When exposed to antifibrotic agent PF3644022, a potent ATP-competitive MK2 inhibitor, pulmonary fibroblasts cultured in IPF matrix scaffolds demonstrated significant reduction in cell number compared to untreated fibroblasts over 6 days. PF3644022 also reduced expression of key IPF-associated genes COL1A1 and ACTA2 by fibroblasts in IPF matrix scaffolds (Fig. 6B) – an expected result not observed in fibroblasts cultured on plastic. Similarly, PF3644022 reduced secretion of bFGF by fibroblasts cultured in IPF matrix scaffolds (Fig. 6C). Interestingly, secretion of TGFβ by fibroblasts exposed to PF3644022 trended upward across all substrates (Fig. 6D). Altogether, these results confirm the activation and diseased phenotype of pulmonary fibroblasts cultured in IPF matrix, and demonstrate the feasibility of testing antifibrotic agents in an *in-vitro* substrate environment with IPF disease-specific features not otherwise present in tissue culture plastic or other conventional drug screening platforms.

**Figure 6.**
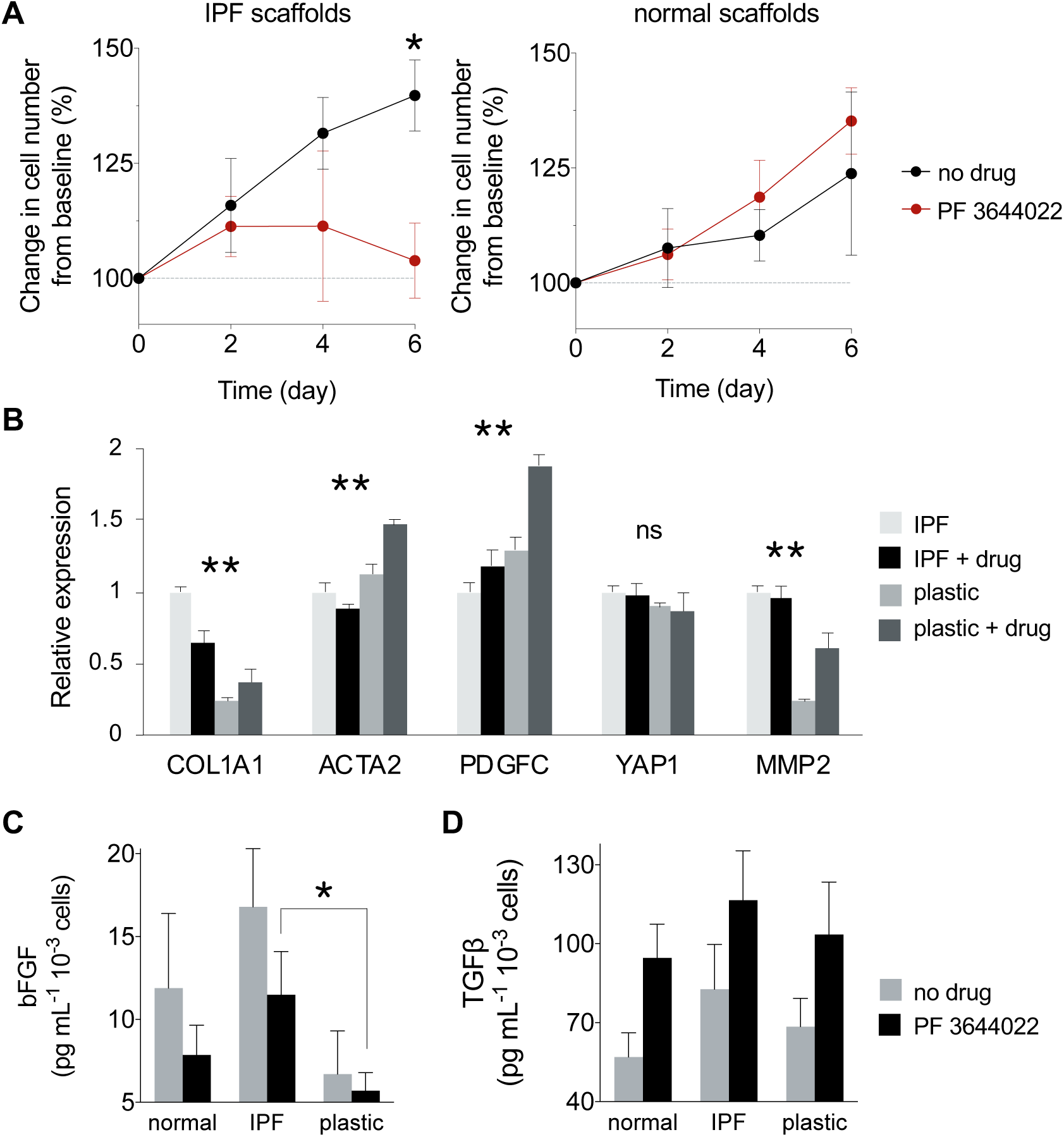
Demonstration of antifibrotic drug testing in IPF lung scaffolds. (**A**) Growth curves of normal human lung fibroblasts over 6 days of treatment with PF3644022. (**B**) Gene expression of normal human lung fibroblasts cultured on IPF scaffolds and tissue culture plastic. * *p* < 0.05, ** *p* < 0.01. Quantification by ELISA of (**C**) basic fibroblast growth factor (bFGF, *p* = 0.0024 by ANOVA) and (**D**) transforming growth factor beta (TGFβ, *p* = 0.0084 by ANOVA) secreted by normal lung fibroblast cultured in IPF and normal lung scaffolds and on tissue culture plastic. All values represent mean ± standard deviation.

## DISCUSSION

Using a physiomimetic approach, we developed an IPF disease-specific 3D cell culture substrate comprised of fibrotic human lung extracellular matrix. Through biomolecular and physico-mechanical characterizations, we show that this disease-specific substrate has numerous physical and compositional features of the human IPF diseased extracellular matrix environment. We also demonstrate the applicability of this substrate for pharmaceutical drug testing. As the critical need for effective IPF drugs persists, human IPF disease-specific cell culture substrates could enable more predictive disease models and drug screening platforms, and accelerate development of new drugs for the treatment of IPF.

Human IPF is a chronic, aging-related disease of unknown etiology typically diagnosed at an advanced stage, and is therefore challenging to model. Both animal^10, 13^ and *in-vitro* models^18, 19, 35–37^ have been used to gain insights into the cellular and molecular mechanisms of IPF. Although animal models of IPF have been developed in mice, rats, hamsters, guinea pigs, rabbits, cats, dogs, sheep, donkeys, horses, and non-human primates,^11, 38–40^ no animal model fully recapitulates the pathophysiology of human IPF – specifically, the histologic pattern of usual interstitial pneumonia and progressive fibrotic disease.^13^ Furthermore, while animal models may inform various aspects of fibrotic lung disease, significant anatomical, biological, and immunological differences from humans reduce pathophysiological relevance to human IPF. Notably, the American Thoracic Society has emphasized the importance of developing ‘humanized’ models of IPF to increase relevance of animal models of IPF to the human disease.^41^

Because animal models of IPF are inherently limited, *in-vitro* models are an indispensable tool in basic and translational studies of human IPF. Previous studies have implicated multiple cellular processes in pulmonary fibrosis including epithelial cell apoptosis^42^, epithelial–mesenchymal transition^43^, and differentiation of fibroblasts to myofibroblasts^44^ that result in significant remodeling and deposition of fibrotic ECM. Conventional two-dimensional (2D) models of IPF typically utilize monolayers of pulmonary myofibroblasts, the primary effector cells of IPF^45^, on tissue culture plastic, enabling mechanistic studies in controlled experimental settings. However, cells cultured in 2D models experience artificial, non-physiological conditions that categorically lack the appropriate three-dimensional (3D) spatial gradients – chemical, mechanical, topographical – in which all lung cells naturally reside within the body. Constrained to one spatial plane, cells in 2D models are immobilized, experience limited cell–cell interactions, and display inhibited cytokinesis and chemotaxis, artificial flattened morphology, unnatural apical-basal polarization, and abnormal integrin and cell-surface receptor expression and distribution.^46–48^ Furthermore, tissue culture plastic has non-physiological topography and stiffness (> 1,000 kPa)^49^, which has been shown to drive atypical cytoskeletal rearrangements^50^, perturb homeostatic gene expression^51^, and induce epigenetic modifications of fibroblasts^52^.

Pulmonary fibroblasts cultured in 3D models, however, adhere to substrates at multiple focal adhesion points^53^, and experience more *in vivo*-like stress–strain^47^ and soluble^54^ gradients. Lung fibroblasts cultured in hydrogels of collagen type I, a structural ECM component upregulated in IPF, previously displayed contraction of collagen hydrogels, whose resistance to cell-generated forces was proportional to expression of αSMA by fibroblasts.^55^ Notably, collagen type I hydrogels are comprised of a single structural ECM component, and thus lack the complex signaling and regulation of the multi-component ECM in the fibrotic disease environment. As paracrine, cell– cell, and cell–matrix interactions are known to drive progression of fibrotic lung disease,^56^ disease-specific 3D cell culture substrates are critical for improving *in-vitro* models of IPF, and should ideally recapitulate the structure, mechanics, and biochemical composition of diseased human lung tissue.

In this study, biochemical and mass spectrometry analyses confirmed that IPF matrix scaffolds had: (*i*) increased collagens and decreased elastin consistent with increased stiffness and decreased compliance, (*ii*) increased proteoglycans, whose covalently bound glycosaminoglycan side chains chondroitin sulfate, dermatan sulfate, heparan sulfate, and hyaluronic acid have been shown to be structurally altered and increased in IPF lungs^57^, and (*iii*) abnormal profile of glycoproteins. Proteoglycans influence viscoelastic properties, cell differentiation, and tissue morphogenesis, and in particular heparan sulfate coordinates ligand–receptor binding of FGF, PDGF, TGFβ, and VEGF^58^ – growth factors involved in pathologic tissue remodeling and detected in IPF matrix scaffolds. Biglycan, a small leucine-rich proteoglycan (SLRP) known to be altered in fibrosis and correlated with lung mechanics through influence on ECM assembly^59^, was increased over 600% in IPF matrix scaffolds. The perturbed profile of glycoproteins in IPF matrix scaffolds included: increased fibrillin 2, a collagenase-resistant glycoprotein that is associated with the 10-nm microfibrils of the basal lamina and regulates the bioavailability of TGBβ through latent transforming growth factor β binding proteins (LTBP)^60^, and increased periostin, a matricellular glycoprotein that promotes fibroblast proliferation, localization of fibrogenic growth factors, collagen type I production, and collagen crosslinking^61^. Notably, the loss of basement membrane components including collagen type IV^29^, laminin, and nidogen in IPF tissues and matrix scaffolds suggests that the use of basement membrane extracts such as Matrigel in models of IPF has minimal pathophysiologic relevance.

The stiffness of fibrotic lung tissue (60 ± 40 kPa) is significantly higher than the stiffness of normal lung tissue (7 ± 6 kPa)^35, 49^, which has critical implications for the stiffness of cell culture substrates in models of IPF, especially for *in-vitro* culture of pulmonary fibroblasts, which exhibit complex mechanotransduction^20, 62^ and have ‘mechanical memory’^63^. In this study, fibrotic human lung ECM scaffolds recapitulated the mechanical differences between normal and fibrotic lung tissues (Fig. 4), and supported increased secretion of bFGF and TGFβ by normal human lung fibroblasts (Fig. 5E,F), suggesting that the IPF matrix scaffolds have disease-specific mechanics and regulatory signals relevant to human IPF. Notably, pulmonary fibroblasts cultured in fibrotic lung ECM previously confirmed the regulatory role of ECM in the activation of myofibroblasts *in vitro*^36^, and demonstrated significant effects of substrate stiffness on fibroblast activation and differentiation into myofibroblasts. Myofibroblast differentiation has also been shown to be driven by increased ECM stiffness through mechanisms independent of TGFβ^20^. Interestingly, previous studies wherein normal and IPF fibroblasts were cultured across ECM from normal and IPF lungs revealed that IPF ECM had a greater influence on fibroblast gene expression than cell origin^64^, further indicating the central role of ECM in regulating disease-associated gene expression. Altogether, these results highlight the critical importance of providing disease-specific signals from the ECM environment in models of fibrotic lung disease.

In spite of decades of basic and translational research, the persisting struggle to successfully translate promising preclinical drug candidates to drugs approved to treat IPF highlights the limited effectiveness of disease models used in IPF drug discovery, which is likely attributable to the failure of IPF models to recapitulate key pathophysiological features of the human disease. Early-stage drug discovery assays are typically conducted on tissue culture plastic (e.g., polystyrene) with or without collagen type I coating, and supplemental TGFβ (e.g., 1 – 5 ng mL^-1^) to activate primary or immortalized human lung fibroblasts – an entrenched *in-vitro* system that has minimal pathophysiological relevance to the human IPF disease setting. With significant financial costs and scientific, medical, and regulatory challenges associated with conducting clinical trials in patients with IPF, preclinical assessments of antifibrotic compounds must be sufficiently robust to inform go/no go decision making and yield reliably predictive data in order to maximize the likelihood of advancing promising drug candidates to clinical trials.

In this study, we demonstrated the use of IPF disease-specific ECM in a 3D cell-based assay of antifibrotic agent PF3644022 (an MK2 inhibitor in IPF model)^65^. As expected, fibroblasts cultured on fibrotic lung ECM scaffolds and treated with PF3644022 exhibited greater sensitivity and drug response, significantly different gene expression, and downregulation of genes associated with ECM production compared to cells cultured on tissue culture plastic. We envision that disease-specific ECM may be applicable across multiple stages of the drug discovery pipeline, from target selection and hit identification through lead identification and optimization. The use of disease-specific ECM substrates is consistent with the set of principles^66^ defined for ‘disease-relevant assays’ that specifically recommend ensuring: (*i*) substrate tension and mechanical forces are appropriate, and (*ii*) extracellular matrix composition is relevant, with appropriate tissue architecture, cell differentiation and function to enhance clinical translation of the *in-vitro* assay. Ultimately, implementation of disease-specific ECM components or substrates into preclinical human disease models and cell-based screening assays could increase clinical relevance and success rates.

There are several limitations to the present study: (*1*) This study investigated a small number of human lungs (*n* = 6 total, *n* = 3 per group). Although this study was conducted with the minimum number of human lungs required to achieve statistical significance between groups, investigation of larger numbers of IPF lungs would offer opportunities for deeper statistical analyses and potential correlations between matrix characteristics and disease phenotypes. (*2*) As IPF lung specimens were procured from explanted tissues following lung transplantation, this study only investigated fibrotic lung matrix from end-stage disease. While diagnosis of IPF remains a significant clinical challenge, procurement of fibrotic human lung ECM at earlier stages of fibrotic disease may not be feasible. (*3*) Human donor tissues present intrinsic biological variability that could confound experimental results. To minimize variability between donors, acceptance criteria for lungs were tightly defined and strictly implemented. Furthermore, as IPF is a disease with demonstrable spatiotemporal heterogeneity, extensive histopathologic review was conducted by a lung transplant pathologist to ensure only tissues and scaffolds with fibrosis scores ≥ 2 were utilized. (*4*) Only one antifibrotic drug was evaluated in this study. Future studies will investigate additional compounds to provide further evidence of the utility and benefits of IPF disease-specific ECM substrates.

The IPF matrix scaffolds developed in this study may be useful for cell-based assays, but may have limited applicability to high throughput drug screening systems, which typically utilize rapid optical readouts in 96-and 384-well plate formats. Therefore, an alternative format of fibrotic lung matrix, e.g., hydrogel, may be more suitable for high throughput applications. Future studies will explore the development of additional IPF disease-specific ECM formats to address broader research and development applications such as ‘IPF-on-chip’. As IPF disease progression is driven by a combination of lung and immune cell–cell and cell–matrix interactions, future studies will also investigate co-cultures with pulmonary macrophages, epithelial and smooth muscle cells, and the effects of IPF-associated growth factors on lung cells *in vitro*. Altogether, an *in-vitro* model with a disease-specific substrate of human IPF disease environment can help elucidate underlying idiopathologies of IPF, enable development of effective IPF therapeutics, and may serve as a template approach for the development of fibrosis-specific cell culture substrates in other human organs and tissues susceptible to fibrotic disease.

## CONCLUSIONS

We developed a pulmonary fibrosis-specific cell culture substrate comprised of intact fibrotic lung extracellular matrix that recapitulated *in vitro* key features of the human IPF disease environment and supported the disease-associated phenotype of human lung fibroblasts. We also demonstrated feasibility of testing antifibrotic agents using this substrate, which may be applicable in cell-based assays in early-stage drug discovery.

## ACKNOWLEDGEMENTS

The authors would like to thank M. Bacchetta and G. Vunjak–Novakovic for discussions of the experimental design and results; M. Bacchetta and Y. Tipograf for coordinating receipt of human lung tissues; Albert Einstein College of Medicine Analytical Imaging Facility for slide scanning; C. Marboe for conducting blinded pathologic assessments; and Weill Cornell Microscopy and Image Analysis Core Facility staff, including L. Cohen-Gould and J. P. Jimenez for transmission electron microscopy imaging services. The authors gratefully acknowledge funding support (1R43HL144341) from the National Institutes of Health.

## AUTHOR CONTRIBUTIONS

I.G., E.A., and J.D.O. designed the study. I.G., E.A., J. X., N. K., A. N., and E.G. performed experiments. I.G., E.A., and J.D.O. co-analyzed data, co-wrote the manuscript.

---

**Supplementary Figure 1.**
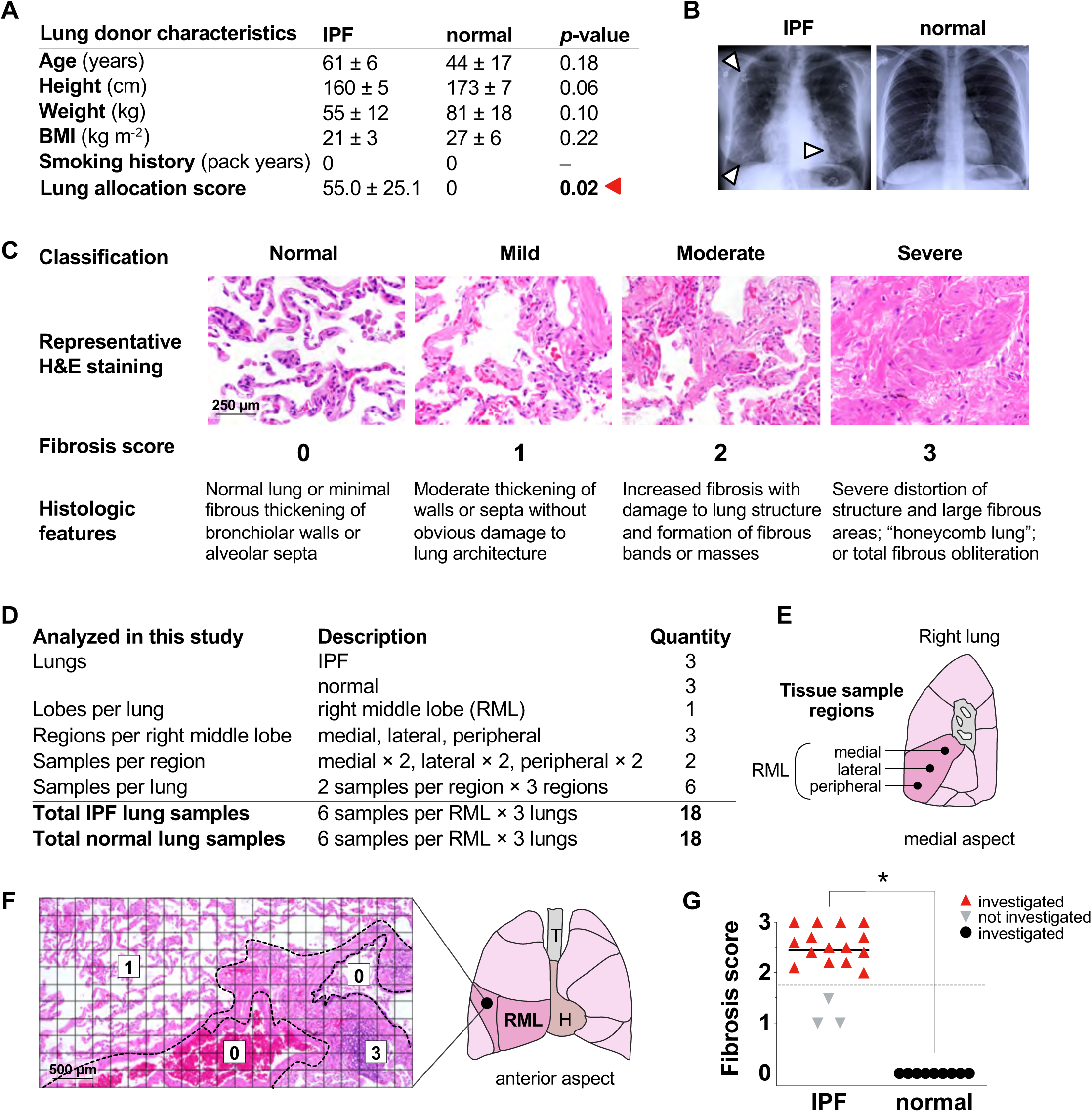
Characterization of human IPF and normal lung tissues. (**A**) Donor characteristics of IPF and normal lung tissues. (**B**) Representative chest radiographs of IPF and normal lung donors. Arrows indicate changed lung shape, decreased lung volume, and increased radiopacity consistent with pulmonary fibrosis. (**C**) Fibrosis scoring rubric used to assess the extent of architectural disruption and fibrosis in human lung tissues. (**D**) Overview of the description and quantity of tissue samples analyzed in this study. Samples were collected from three regions (i.e., medial, lateral, peripheral) of right middle lobes. (**F**) Demonstration of quantitative image analysis method used for fibrosis scoring. For each fibrosis score, a weighted average is calculated from the ratios of the total area. Five high-power fields were analyzed per region, and an average fibrosis score was calculated for each region. (**G**) Fibrosis scores of all regions of IPF and normal lungs investigated in this study. Only regions of IPF lungs with fibrosis score ≥ 2 (red triangle) were investigated in this study. * *p* < 0.001.

**Supplementary Figure 2.**
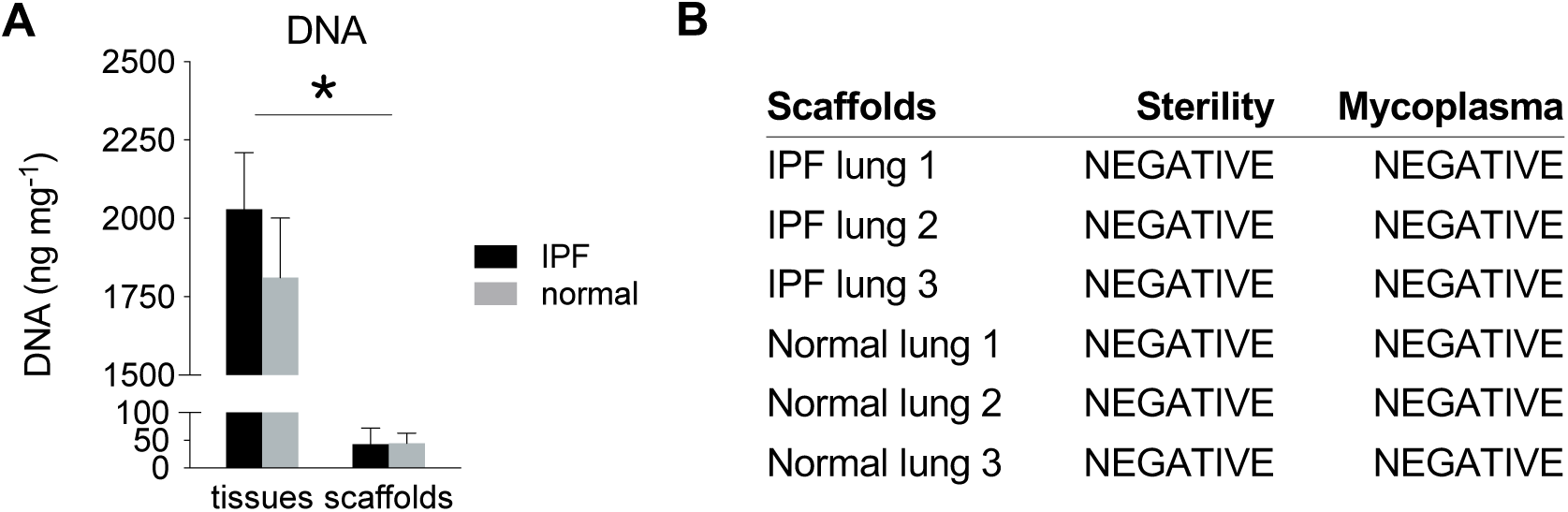
Quality control assays of IPF and normal lung matrix scaffolds. Quantification of (**A**) DNA to confirm removal of nuclear material from IPF and normal lung matrix scaffolds. (**B**) Results of sterility and mycoplasma assays. Prior to use in studies, IPF and normal lung matrix scaffolds were tested for absence of bacteria and fungi. Scaffolds were also tested for absence of mycoplasma using MycoAlert PLUS Mycoplasma Detection Assay. * *p* < 0.001.

**Supplementary Table 1.** Antibodies and ELISA kits.

| Antibody | Application | Vendor | Product number | Dilution |
| --- | --- | --- | --- | --- |
| Rabbit anti-alpha smooth muscle actin | IHC-P | Cell Signaling Technology | 19245 | 1 : 500 |
| Rabbit anti-fibrillin 2 | IHC-P | Sigma Life Science | HPA012853 | 1 : 50 |
| Rabbit anti-Ki67 | IHC-P | ThermoFisher Scientific | PA1-38032 | 1 : 500 |
| Rabbit anti-laminin γ1 | IHC-P | Abcam | ab233389 | 1 : 2000 |
| Rabbit anti-matrix gla protein | IHC-P | LS Bio | LS-B14824 | 1 : 100 |
| Rabbit anti-periostin | IHC-P | Abcam | ab14041 | 1 : 300 |

**Supplementary Table 1 | Antibodies and ELISA kits.**
| ELISA kit | Application | Vendor | Product number |
| --- | --- | --- | --- |
| Human basic fibroblast growth factor | Cell culture media | R&D Systems | DFB50 |
| Human transforming growth factor $\beta$ | Cell culture media | R&D Systems | DB100B |

**Supplementary Table 2.**
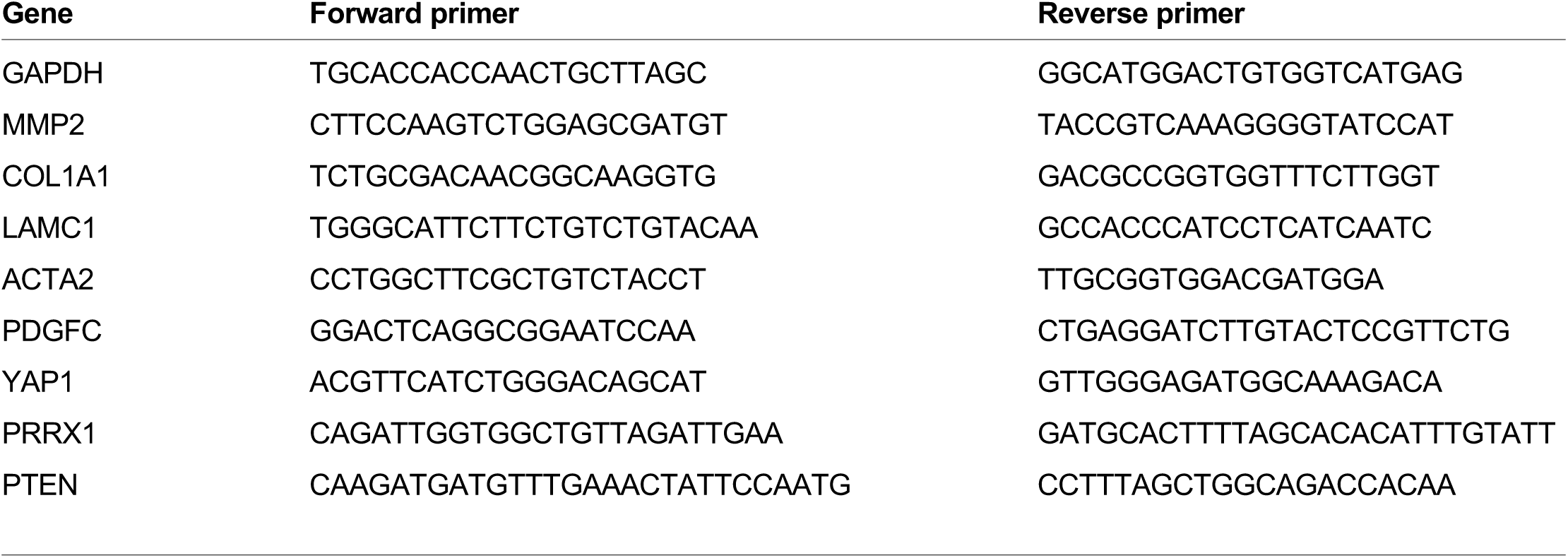
Primers.

**Supplementary Table 3.**
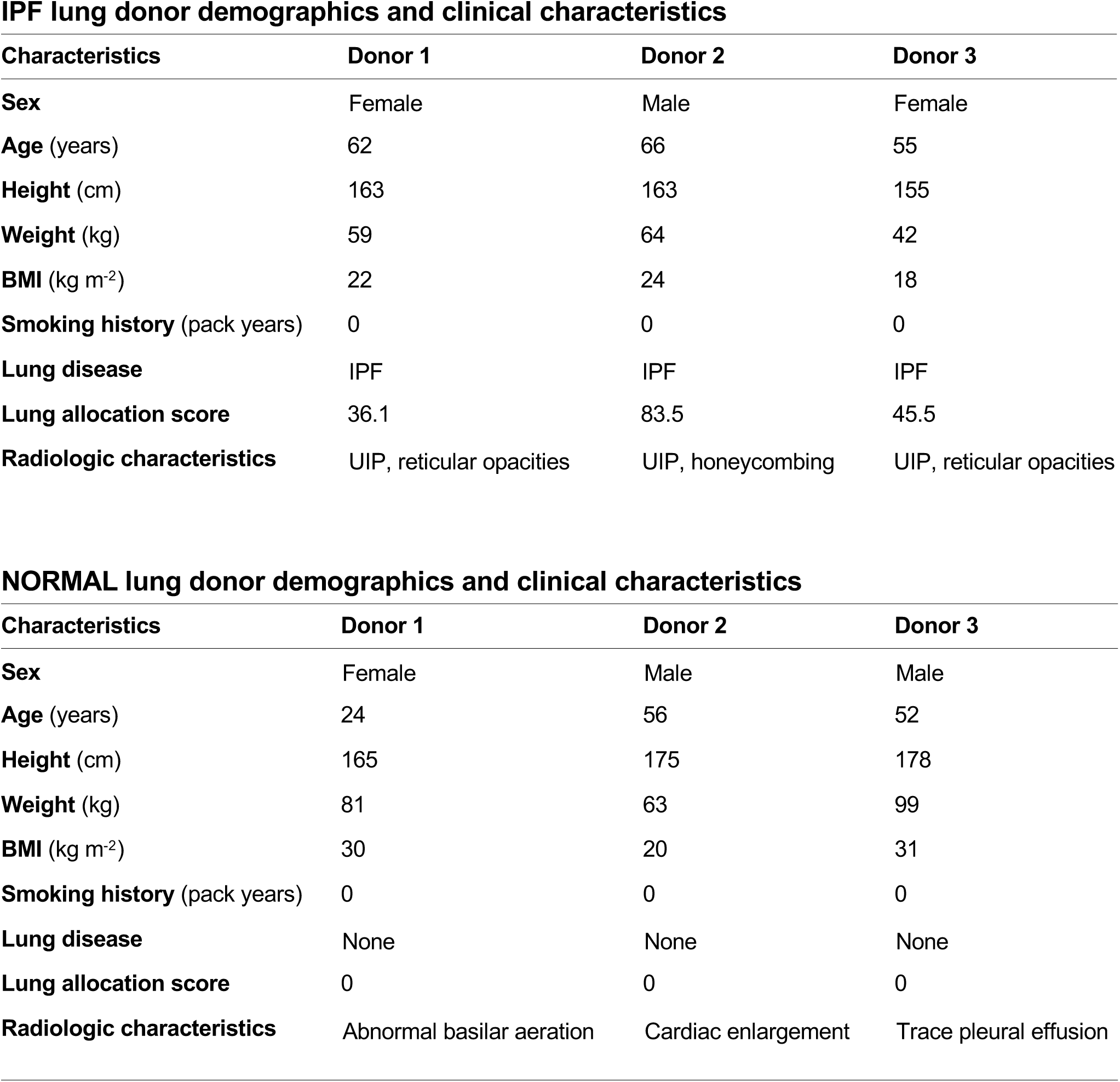
Demographic and clinical characteristics of human lung donors. UIP, usual interstitial pneumonia.

**IPF lung donor demographics and clinical characteristics**
| Characteristics | Donor 1 | Donor 2 | Donor 3 |
| --- | --- | --- | --- |
| **Sex** | Female | Male | Female |
| **Age** (years) | 62 | 66 | 55 |
| **Height** (cm) | 163 | 163 | 155 |
| **Weight** (kg) | 59 | 64 | 42 |
| **BMI** (kg m<sup>-2</sup>) | 22 | 24 | 18 |
| **Smoking history** (pack years) | 0 | 0 | 0 |
| **Lung disease** | IPF | IPF | IPF |
| **Lung allocation score** | 36.1 | 83.5 | 45.5 |
| **Radiologic characteristics** | UIP, reticular opacities | UIP, honeycombing | UIP, reticular opacities |

**Supplementary Table 3 | Demographics and clinical characteristics of human lung donors.** UIP, usual interstitial pneumonia.
| Characteristics | Donor 1 | Donor 2 | Donor 3 |
| --- | --- | --- | --- |
| **Sex** | Female | Male | Male |
| **Age** (years) | 24 | 56 | 52 |
| **Height** (cm) | 165 | 175 | 178 |
| **Weight** (kg) | 81 | 63 | 99 |
| **BMI** (kg m<sup>-2</sup>) | 30 | 20 | 31 |
| **Smoking history** (pack years) | 0 | 0 | 0 |
| **Lung disease** | None | None | None |
| **Lung allocation score** | 0 | 0 | 0 |
| **Radiologic characteristics** | Abnormal basilar aeration | Cardiac enlargement | Trace pleural effusion |

**Supplementary Table 4.** Quantification of growth factors in IPF and normal lung tissues. Growth factor concentrations were measured by multiplex growth factor array. Green arrow (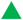) indicates positive fold change (increase) from normal in concentration of growth factors. Red arrow (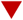) indicates negative fold change (decrease) from normal in concentration of growth factors. ND: not detected. * IPF-specific growth factor not detected in normal lung tissue.

| Growth factor | Description | Concentration (pg mL<sup>-1</sup>) |  | Fold change from normal |
| --- | --- | --- | --- | --- |
|  |  | Normal tissue | IPF tissue |  |
| TGF-β3 | Transforming growth factor β3 | ND | 65.8 | * |
| HB-EGF | Heparin-binding EGF-like growth factor | ND | 4.2 | * |
| IGFBP-1 | Insulin-like growth factor binding protein 1 | 0.5 | 86.8 | 159.5 ▲ |
| bFGF | Basic fibroblast growth factor | 10.1 | 213.8 | 21.2 ▲ |
| EG-VEGF | Endocrine gland-derived vascular endothelial growth factor | 2.2 | 38.1 | 17.3 ▲ |
| BDNF | Brain-derived neurotrophic factor | 31.1 | 145.4 | 4.7 ▲ |
| GDF-15 | Growth differentiation factor 15 | 97.8 | 243.4 | 2.5 ▲ |
| PDGF-AA | Platelet-derived growth factor AA | 228.1 | 486.7 | 2.1 ▲ |
| IGFBP-6 | Insulin-like growth factor binding protein 6 | 123.0 | 228.5 | 1.9 ▲ |
| HGF | Hepatocyte growth factor | 13313.2 | 19831.9 | 1.5 ▲ |
| VEGF | Vascular endothelial growth factor | 135.9 | 158.8 | 1.2 ▲ |
| EGF R | Epidermal growth factor receptor | 17343.2 | 12828.8 | 0.7 ▼ |
| OPG | Osteoprotegerin | 60.8 | 33.8 | 0.6 ▼ |

## SUPPLEMENTARY INFORMATION

### Characterization of human IPF and normal lung tissues

Donor characteristics of IPF and normal lung tissues were analyzed to confirm that there were no significant differences in age, height, weight, body mass index, or smoking history (Supplementary Fig. 1A, Supplementary Table 3). The mean lung allocation score for IPF donors was 55.0 ± 25.1, which was the only significant difference between IPF and normal lung donors (*p* < 0.05). Chest radiographs of lung donors confirmed absence of apparent injury or underlying disease in normal lungs, and enabled assessment of the extent and distribution of pulmonary fibrosis in IPF lungs. In contrast to normal lungs, which appeared radiolucent and aerated, IPF lungs displayed a morphologic pattern of usual interstitial pneumonia consistent with IPF and marked by diffuse radiopacities, reticulation, architectural distortion, and honeycombing, especially in peripheral and basal regions (Supplementary Fig. 1B).

In order to characterize the histopathology of all tissues investigated, a fibrosis scoring rubric was used to assess the extent of architectural disruption and fibrosis (Supplementary Fig. 1C). Samples were systematically collected from the medial and lateral regions of the right middle and right lower lobes. Histologic samples were evaluated by light microscopy, and fibrosis scores were assigned and averaged across five high-power fields. All high-power fields were subjected to imaging analyses to quantify the relative areas corresponding to each fibrosis score (Supplementary Fig. 1F). To ensure to the maximum possible extent a consistent degree of pulmonary fibrosis across all samples, only samples with average fibrosis score ≥ 2 (moderate or severe fibrosis) were investigated in this study (Supplementary Fig. 1G).

### Mass spectrometry

Protein profiling included short gel SDS-PAGE, in-gel digestion with trypsin, and 2 hours LC-MS/MS. Samples were weighed and suspended in 130 µL of 2.0% modified RIPA buffer with 1.6 mm stainless steel beads. Samples were homogenized in a Next Advance Bullet Blender for 3 minutes at speed 10, then heated at 100°C for 30 minutes. Samples were then sonicated and clarified by centrifugation. The protein concentration of the extract was determined using Qubit fluorometry (Life Technologies). Each sample (10 μg) was processed by 2 cm SDS-PAGE using a 10% Bis-Tris NuPAGE Novex mini gel (ThermoFisher) with the MES buffer system. The mobility region was excised and processed by in-gel digestion with trypsin using a ProGest robot (DigiLab) with the following protocol: (1) Wash with 25 mM ammonium bicarbonate followed by acetonitrile. (2) Reduce with 10 mM dithiothreitol at 60°C followed by alkylation with 50 mM iodoacetamide at room temperature. (3) Digest with sequencing grade trypsin (Promega) at 37°C for 4 hours. (4) Quench with formic acid and analyzed without further processing.

Half of each digest was analyzed by nano LC-MS/MS with a Waters NanoAcquity HPLC system interfaced to a mass spectrometer (Fusion Lumos, ThermoFisher). Peptides were loaded on a trapping column and eluted over a 75 μm analytical column at 350 nL min^-1^ with a reverse phase gradient for 2 hours. Both columns were packed with Luna C18 resin (Phenomenex). The mass spectrometer was operated in data-dependent mode, with the Orbitrap operating at 60,000 FWHM and 15,000 FWHM for MS and MS/MS, respectively. The instrument was run with a 3 second cycle for MS and MS/MS. Advanced Precursor Determination was employed. Data were searched using a local copy of Mascot (Matrix Science) with the following parameters: Enzyme: Trypsin/P; Database: SwissProt Human (concatenated forward and reverse plus common contaminants); Fixed modification: Carbamidomethyl (C); Variable modifications: Oxidation (M/P), Acetyl (N-term), Pyro-Glu (N-term Q), Deamidation (N,Q); Mass values: Monoisotopic; Peptide Mass Tolerance: 10 ppm; Fragment Mass Tolerance: 0.02 Da; Max Missed Cleavages: 2. Mascot DAT files were parsed into Scaffold (Proteome software) for validation and filtering to create a non-redundant list per sample. Data were filtered using at least 1% protein and peptide FDR, and requiring at least two unique peptides per protein.

